# Sex-dependent Differences in the Genomic Profile of Lingual Sensory Neurons in Naïve and Tongue-Tumor Bearing Mice

**DOI:** 10.1101/2023.01.14.524011

**Authors:** Tarek Ibrahim, Ping Wu, Li-Ju Wang, Chang Fang-Mei, Josue Murillo, Jaclyn Merlo, Alexei Tumanov, Zhao Lai, Korri Weldon, Yidong Chen, Shivani Ruparel

**Affiliations:** Department of Endodontics, School of Dentistry, University of Texas Health San Antonio, USA; Greehey Children’s Cancer Institute, University of Texas Health San Antonio, USA; Department of Population Health Sciences, University of Texas Health at San Antonio, USA; Department of Microbiology, Immunology and Molecular Genetics, University of Texas Health San Antonio, USA; Department of Molecular Medicine, University of Texas Health at San Antonio, San Antonio, TX, USA

## Abstract

Mechanisms of sex-dependent orofacial pain are widely understudied. A significant gap in knowledge exists about comprehensive regulation of tissue-specific trigeminal sensory neurons in diseased state of both sexes. Using RNA sequencing of FACS sorted retro-labeled sensory neurons innervating tongue tissue, we determined changes in transcriptomic profiles in males and female mice under naïve as well as tongue-tumor bearing conditions Our data revealed the following interesting findings: 1) Tongue tissue of female mice was innervated with higher number of trigeminal neurons compared to males; 2) Naïve female neurons innervating the tongue exclusively expressed immune cell markers such as Csf1R, C1qa and others, that weren’t expressed in males. This was validated by Immunohistochemistry. 4) Accordingly, immune cell markers such as Csf1 exclusively sensitized TRPV1 responses in female TG neurons. 3) Male neurons were more tightly regulated than female neurons upon tumor growth and very few differentially expressed genes (DEGs) overlapped between the sexes, 5) Male DEGs contained higher number of transcription factors whereas female DEGs contained higher number of enzymes, cytokines and chemokines. Collectively, this is the first study to characterize the effect of sex as well as of tongue-tumor on global gene expression, pathways and molecular function of tongue-innervating sensory neurons.

## Introduction

Sex-dependent differences in orofacial pain has been clinically well-established with higher prevalence in women in several different pathological conditions^1-6^; although the mechanism of sex-differences remain elusive. It is known that sensory neurons including trigeminal neurons, that regulate pain in the orofacial region, are genomically different in males and females even under naïve conditions ^7-10^. These studies have primarily been conducted using the entire neuronal population of the ganglionic tissue. However, several reports reveal that sensory innervation can be different with each tissue type ^7,11-14^. Accordingly, we have previously identified subsets of sensory neurons expressed in mouse tongue^13^ that are varied from those innervating the masseter muscle^15^. Therefore, it is vital in delineating tissue-specific sex-dependent differences in trigeminal sensory neurons. Moreover, a significant gap-in knowledge exists for sex-specific changes of trigeminal neurons that specifically innervate diseased-tissues. Such studies can provide crucial information about the regulation of trigeminal sensory neurons in tissue-specific pathologies.

The tongue is among the vital organs of the orofacial region as it controls many essential daily activities such as speech, licking, taste, chewing and swallowing. Importantly, many lingual ailments cause acute and chronic pain leading to very distressing and debilitating quality of life ^16-29^. In fact, among all chronic orofacial pain conditions that pose a challenge in management, three are known to primarily affect the tongue. These include tongue cancer, oral mucositis and burning mouth syndrome^23-26,30-42^. Interestingly, sex-dependent differences in the manifestation of pain have been reported for each of these conditions^30,31,35,36,43-48^.

Therefore, the current study identified the differences in tongue-innervating sensory neurons between males and females under naïve and diseased state. We used tongue cancer as our disease model as approximately 50% of oral cancer patients report pain throughout the course of the disease and of these, the prevalence of pain is highest in tongue cancer patients^23-26 49^. Pain from tongue cancer is extremely weakening and significantly deteriorates patient quality of life in addition to having cancer due to very limited treatment options available. Therefore, using the orthotopic tongue cancer xenograft model in mice, we performed bulk-RNA sequencing of isolated tongue-innervated sensory neurons from males and females, to identify changes in genes, biological processes and molecular function between sexes under normal and tumor-bearing conditions.

## Results

### Isolation of Lingual TG neurons revealed differences in number of innervating neurons between males and females

To assess transcriptomic changes in tongue-innervating TG neurons in males and females under naïve and tumor-bearing conditions, we performed bulk-RNA sequencing of lingual TG neurons. To achieve this, we first isolated retro-labeled TG neurons innervating mouse tongue using WGA-488 and subsequently flow-sorted fluorescently labeled neurons from four groups: Male normal (MN), Male tumor (MT), Female Normal (FN) and Female Tumor (FT). Representative gating strategy for the sorting protocol is shown in **Fig 1A**. Flow sorting obtained an average of approximately 10K to 23K neurons per sample for all four groups (**Fig 1B and Suppl Table 1**). Surprisingly, we found that the number of neurons isolated from all of female samples from both groups (i.e. FN and FT,) were higher (approx. 22000 cells) compared to male groups (i.e MN and MT, approx. 12000 cells) (**Fig 1B**). In potting the percentage of WGA+ neurons sorted over all live events in the samples, we found that percentage of WGA+ neurons isolated from female samples were significantly higher (approx. 3 fold, one-way ANOVA, p<0.05) than in males (**Fig 1C**) suggesting that female mice may have increased number of sensory neurons innervating the tongue tissue compared to males in mice. To further investigate this finding, we employed immunohistochemistry to evaluate nociceptive and non-nociceptive WGA+ neurons in naïve male and female mice. We used TRPV1 to distinguish between the two neuronal classes and found that males had a higher percentage of TRPV1+ tongue innervating neurons compared to females **Fig 1D and E**) (27.6% males vs 19.31% females, two-way ANOVA, p=0.0077). Accordingly, females had a higher percentage of TRPV1 negative tongue innervating neurons than males (Fig 1D and E) (72.38% males vs 80.6% females, two-way ANOVA, p=0.0077).

**Fig 1.**
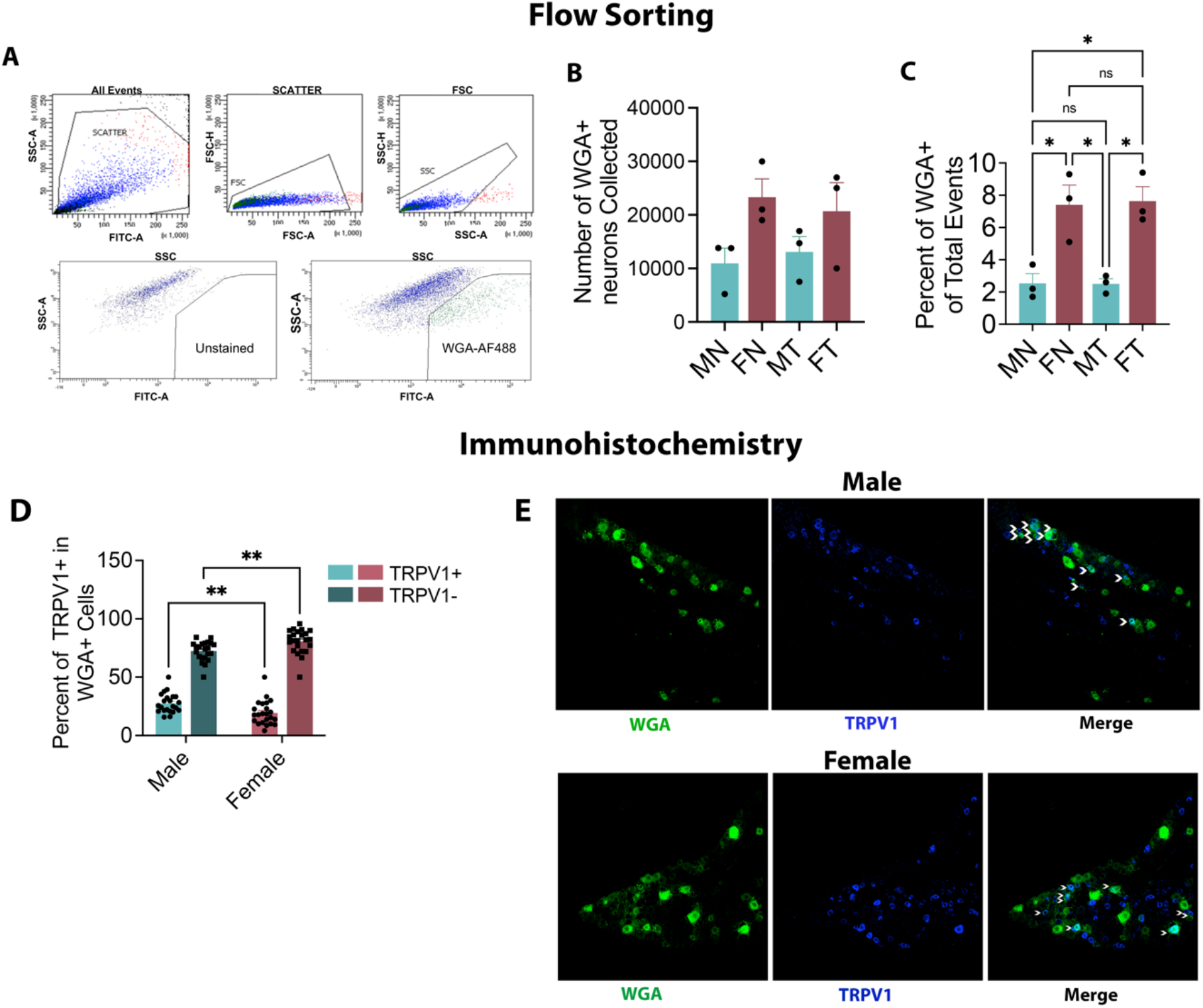
Isolation and Estimation of Tongue-Innervating Sensory Neurons. **A**,**B and C**. Male and Female mice were injected with 3 x 10^5 HSC3 cells in the tongue and at day 13 post-cell inoculation, tongue tissues were bilaterally injected with 1% WGA-488. Normal group received no HSC3 cells. Two days later, TG tissues were dissected to make single-cell suspension and subjected to flow sorting. Animals were grouped into male normal (MN), male tumor (MT), female normal (FN) and female tumor (FT). N=3 samples per group. **A**. Gating Strategy for flow sorting is shown. **B**. Number of cells sorted for each group. Data points represent numbers of cells in each sample. **C**. Percentage of WGA+ neurons of total events in each group. Data points represent percentage of cells in each sample. Data are represented as mean ± SEM and analyzed by one-way ANOVA with Sidak’s post-hoc test.p<0.05. **D and E**. Naïve male and female mice were injected with 1%WGA-488 and 2 days later, TG tissues were harvested for immunohistochemistry. Images were taken at 20x magnification using the Nikon C1 confocal microscope. N=2 per group. **D**. Percentage of TRPV1+/WGA+ and TRPV1-/WGA+ TG neurons in naïve males and females. Each data point represents percentage of neurons in each image. Data are represented as mean ± SEM and analyzed by one-way ANOVA with Sidak’s post-hoc test. **E**. Representative images of WGA and TRPV1 staining of TG tissues in male and females are shown. Arrows indicate colocalization of TRPV1 and WGA.

### Sex-dependent differences in gene expression in lingual neurons of naïve mice

Bulk-RNA sequencing was conducted of flow-sorted WGA+ TG neurons from all four groups: MN, MT, FN and FT. Data were analyzed to identify differentially expressed genes (DEGs) using the criteria: RPKM>5, FC> 1.5 and p<0.05. In comparing FN vs MN groups, we identified a total of 81 DEGs between both sexes after excluding all sex-linked genes. Of these, 30 genes were exclusively expressed in females and only 2 genes exclusively expressed in males (**Fig 2A and B**.). The remaining 49 genes, while expressed in both sexes, were significantly upregulated in females compared to males (**Fig 2C and D**). Details of RPKM and FC of the top 10 genes is listed in **Supplementary Table 2**. Two genes, that were higher in males than females were Fam23a(transmembrane protein 236) and Ddx3y (DEAD box helicase). In further assessing the expression of female-specific genes, we tested the expression of selected two genes: Csf1R and C1qa using immunohistochemistry and confirmed their higher expression in females than in males. Csf1R was expressed in ∼ 42% of all WGA+ neurons in females compared to 5.5% of WGA+neurons in males (Unpaired Student T test, p<0.0001) **(Fig 2E**). Of all Csf1R positive neurons innervating the tongue in females, 18% were found to be TRPV1 positive (**Fig 2F and G**) (Paired Student’s T test, p=0.0049). Similarly, ∼53% of all tongue innervating neurons expressed C1qa in females whereas almost no expression of this gene was found in males (**Fig 2H**) (Unpaired Student T test, p<0.0001). Majority of C1qa positive neurons in the tongue of females were TRPV1 negative (94%) with only a very small proportion of C1qa expression in TRPV1+ nociceptors (6%) (**Fig 2I and J**) (Paired Student’s T Test, p<0.001).

**Fig 2.**
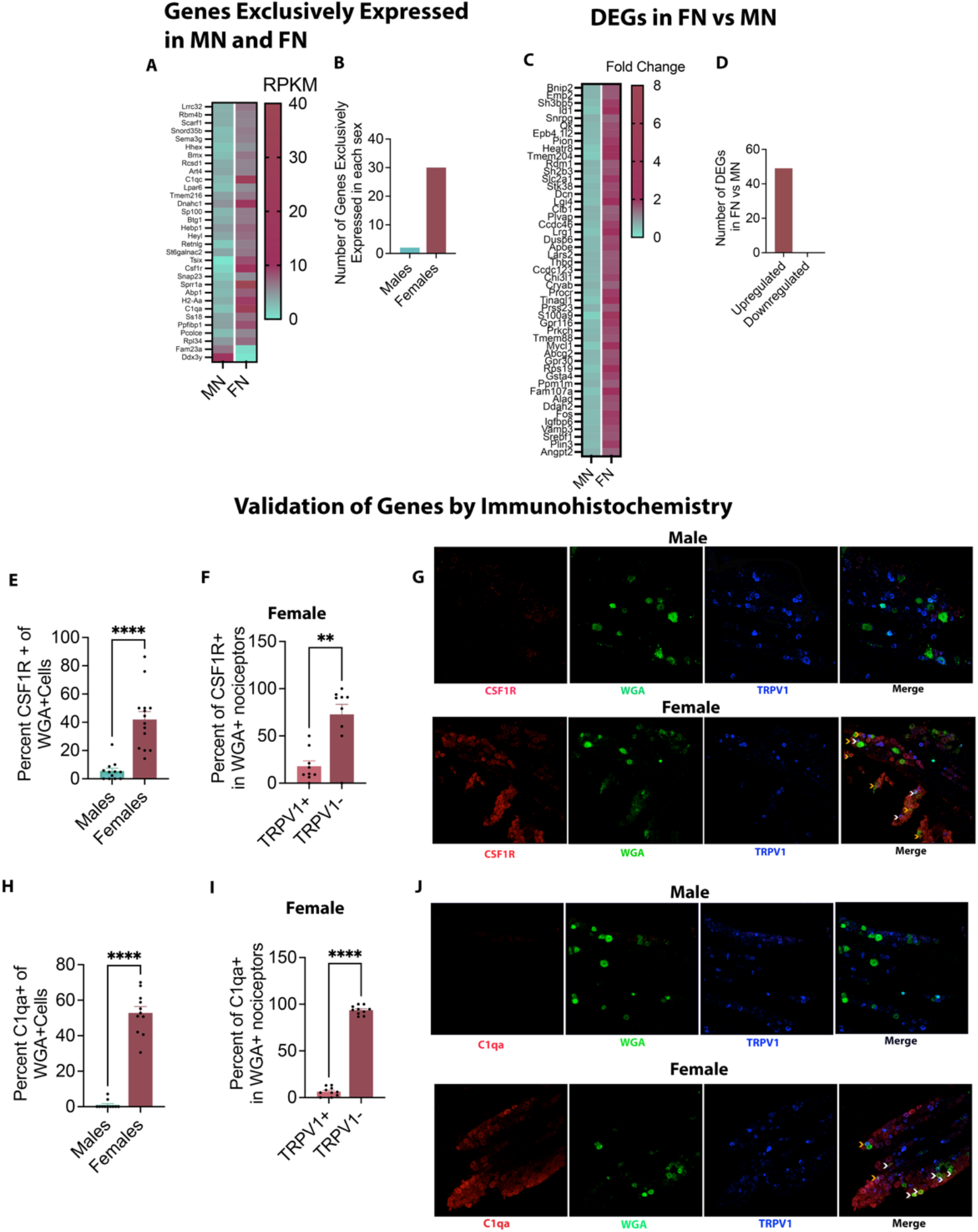
Differentially Expressed Genes in Male versus Female Lingual Neurons. **A-D**. Flow sorted neurons were subjected to RNA sequencing. DEGs were identified in normal male and female mice based on RPKM>5, FC>1.5 and p<0.05. N=3 per group. **A**. Heatmap of genes that were expressed exclusively in tongue-innervating neurons of normal male and female mice. **B**. Number of genes in males and females is depicted as bar graphs. **C**. Heatmap of genes differentially expressed in female normal (FN) vs male normal (MN). **D**. Bar graphs shows number of genes upregulated and downregulated in FN vs MN. **E-J**. Validation of RNA sequencing data in normal male and female by Immunohistochemistry. N=2 per group. Images taken with C1 Nikon Confocal Microscope at 20x magnification. **E**. Percentage of WGA+neurons expressing Csf1R in males and females. Each data point represents percentage of neurons in each image. Data are represented as mean ± SEM and analyzed by unpaired Student’s T Test at p<0.05. **F**. Percentage of Csf1R in lingual TRPV1+ and TRPV1-neurons. Each data point represents percentage of neurons in each image. Data are represented as mean ± SEM and analyzed by paired Student’s T Test at p<0.05 **G**. Representative images of immunostaining of Csf1R and TRPV1 in WGA+ neurons in males and females. White arrows indicate colocalization of Csf1R and WGA whereas orange arrows indicate colocalization of Csf1R, WGA and TRPV1. **H**. Percentage of WGA+neurons expressing C1qa in males and females. Each data point represents percentage of neurons in each image. Data are represented as mean ± SEM and analyzed by unpaired Student’s T Test at p<0.05 **F**. Percentage of C1qa in lingual TRPV1+ and TRPV1-neurons. Each data point represents percentage of neurons in each image. Data are represented as mean ± SEM and analyzed by paired Student’s T Test at p<0.05 **G**. Representative images of immunostaining of C1qa and TRPV1 in WGA+ neurons in males and females. White arrows indicate colocalization of C1qa and WGA whereas orange arrows indicate colocalization of C1qa, WGA and TRPV1.

To assess whether the genes exclusively expressed in females may have a role in nociception, we tested the function of Csf1R in sensitizing nociceptors due to its expression in TRPV1+ neurons. Accordingly, we explored whether its ligand Csf1 sex-selectively increases TRPV1 responses in TG neurons. As shown in **Fig.3A**, proportion of WGA+ and WGA-neurons that were responsive to CAP did not change with Csf1 treatment in females. However, Csf1 significantly increased CAP-evoked calcium accumulation of WGA+ as well as WGA-cells (**Fig3B and 3C**). Expectedly, Csf1 treatment neither altered the percentage of CAP responsive neurons (**Fig.3D**) nor CAP-evoked calcium accumulation in male neurons (**Fig 3E and 3F**). These data indicate that Csf1R in sensory neurons may sex-specifically contribute to nociceptive responses.

**Fig 3.**
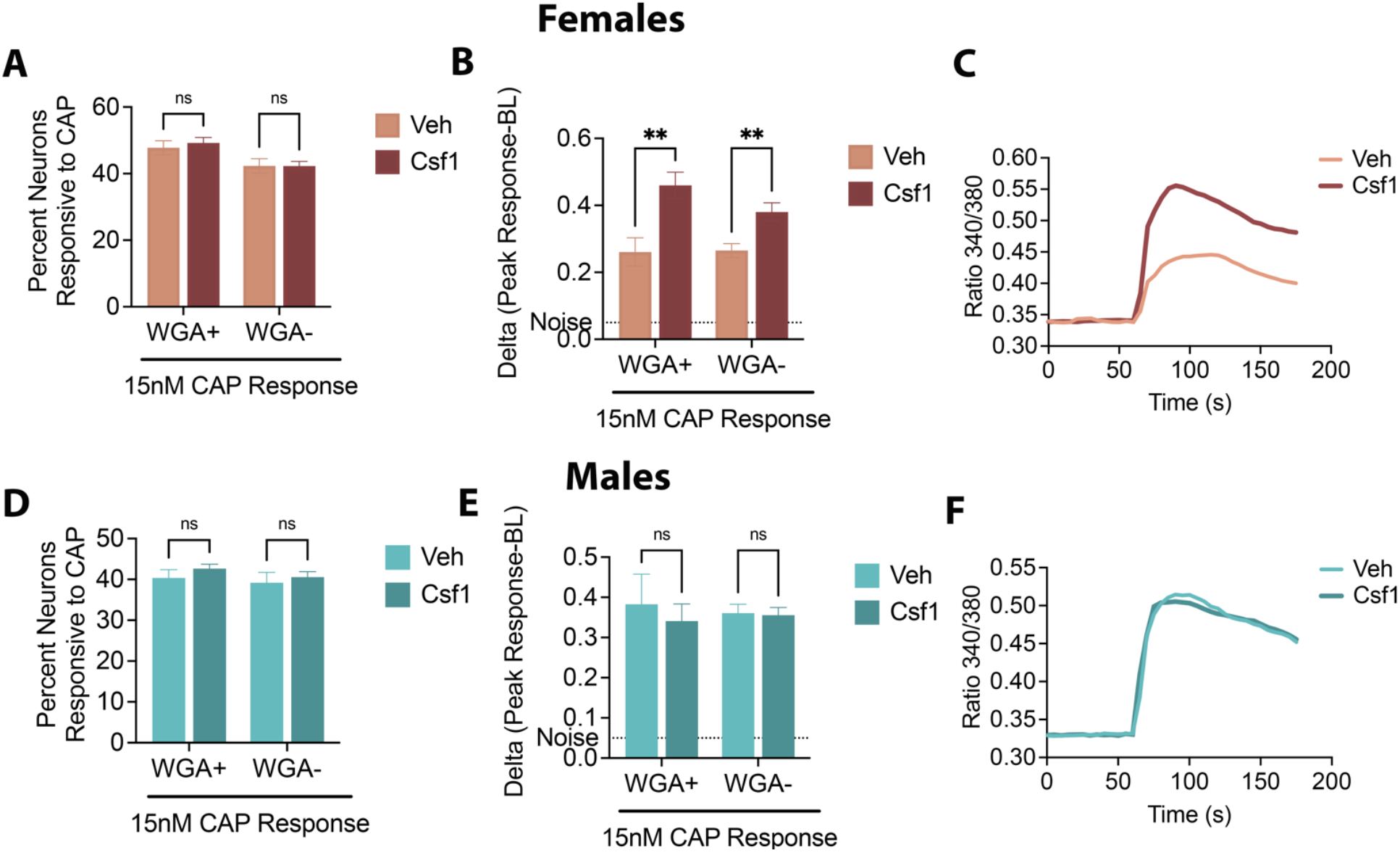
Effect of Csf1 in CAP-evoked calcium influx in male and female TG Neurons. **A, B and C** Naïve female and **D, E and F** naïve male mice were injected with 0.5% WGA and 2 days later, TG neuronal cultures were treated with either vehicle (Veh) or 100ng/ml Csf1 overnight. Following treatment, neurons were recorded for 15nM CAP-evoked calcium response. **A and D**. Data are presented as mean ± SEM of percent neurons responsive to CAP in WGA+ and WGA-neurons. Data analyzed by two-way ANOVA with Sidak post-hoc test at p<0.05 **B and E**. Data are presented as mean ± SEM of delta values calculated by subtracting baseline (BL) from peak CAP response in WGA+ and WGA-neurons. Data analyzed by two-way ANOVA with Sidak post-hoc test at p<0.05. **C and F**. Representative traces of CAP response in Veh and CSF1 treated neurons is shown. Experiments were performed twice on two different days for each sex.

### Changes in Transcriptomic profile of lingual neurons upon tongue tumor growth in males and females

We next compared MT vs MN to elucidate the changes in transcriptome of tongue-innervating neurons upon HSC3-induced tongue tumor in male mice. As shown in **Fig 4A and B**, we observed 83 DEGs, of which 75 genes were upregulated and 8 genes were down-regulated in tumor-bearing group compared to control. Of the 75 up-regulated genes, two were sex-linked genes (**Fig 4B**) (i.e Timp1 and Rbm3). The Top 3 genes that were upregulated upon tumor growth were found to be Sprr1a (small proline-rich protein 1A, FC=19.09, p<0.01), Gal (Galanin, FC=5.11, p<0.0001) and FGF23 (fibroblast growth factor receptor 23, FC=4.22, p<0.001) (**Fig 4C**). The top three genes downregulated in the tumor-bearing group of male mice were Gm2058 (ubiquitin-conjugating enzyme E2H pseudogene, FC=0.211, p<0.05), Sst (somatostatin, FC=0.313, p<0.001), Ms4a3 (membrane-spanning 4-domains, subfamily A, member 3, FC=0.327, p<0.001) (**Fig 4C**).

**Fig 4.**
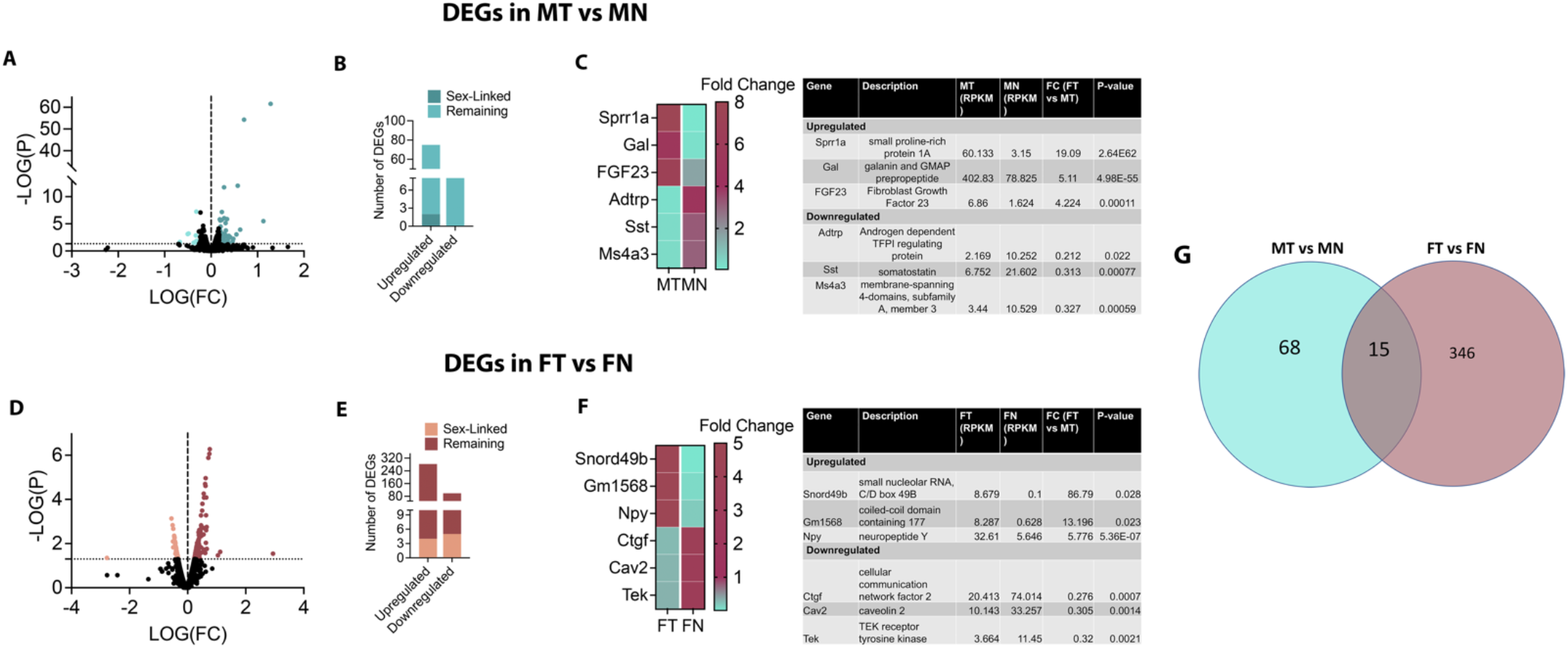
Effect of tongue tumor on transcriptomic profile of lingual neurons in male and female mice. DEGs were identified by conducting two comparisons. **A**. Volcanic Plots for all genes for MT vs MN comparison. DEGs identified are colored showing downregulated genes on the left and upregulated genes on the right. **B**. Number of upregulated and downregulated genes are plotted as bar graph. **C**. Top three upregulated and downregulated genes in MT vs MN are plotted as heatmap as well as tabulated for values of RPKM, Fold change (FC) and p-value. Data for heatmap plotted as fold change. Similarly, **D**. Volcanic plots for FT vs FN. **E**. Bar graph for upregulated and downregulated genes for FT vs FN. **F**. Top 3 three upregulated and downregulated DEGs in FT vs FN as heatmaps and tabulated for RPKM, FC and p-values. Data in heatmap plotted as fold change. **G**. Venn Diagram indicating number of overlapping and non-overlapping between MT vs MN and FT vs FN.

Unlike in males, the number of DEGs in females upon tumor growth were considerably higher (**Fig 4D**). Comparing FT vs FN, we observed a total of 382 DEGs (**Fig 4D and E**), out of which 283 genes were upregulated and 99 genes were downregulated. Four upregulated (i.e Timp1, Arxes1, Timm17b and Rnf113a1) and five downregulated (i.e. Bmx, Gyk, Map3k15, Tmem47 and Slitrk4) genes were sex-linked (**Fig 4E**). The top three upregulated genes in FT vs FN comparison were found to be Snord49b (small nucleolar RNA C/D box 49B, FC =86.79, p=0.028), Gm1568 (coiled-coil domain containing 177, FC=13.196, p=0.0023) and Npy (neuropeptide Y, FC=5.776, p<0.0001) (**Fig 4F**). The top three downregulated genes in tumor-bearing group of female mice were Ctgf (cellular communication network factor 2, FC=0.276, p=0.0007), Cav2 (caveolin 2, FC=0.305, p<0.0014) and Tek (TEK receptor tyrosine kinase, FC=0.32, p<0.0021) (**Fig 4F**).

Between DEGs observed from MT vs MN and FT vs FN, we found 18 genes that were common to both comparisons whereas majority of the genes were regulated in a sex-specific manner (**Fig 4G**). Interestingly though, out of the 18 common DEGs, 11 were upregulated in both comparisons whereas 3 genes that were upregulated in MT vs MN were significantly downregulated in FT vs FN (Suppl Table 3). In contrast 4 genes that were downregulated in MT vs MN were upregulated in FT vs FN (**Suppl Table 3**).

Collectively, these data suggests that the regulation of lingual neurons by the tongue tumor may be considerably different in males and females.

### Biological Processes and Molecular Function Regulated Upon Tumor Growth in Males and Females

We next conducted gene ontology analyses to identify key biological processes in males and females upon tumor growth. Fourteen distinct biological processes were identified with upregulated DEGs in males upon tumor growth (**Fig 5A**). These processes pertained to signaling mechanisms, immune process and inflammation, metabolic processes as well as some others including response to glucocorticoid stimulus, circadian regulation glial cell proliferation and apoptosis. Majority of DEGs in males were involved in signaling pathways and immune and inflammatory processes with predominant processes being second messenger signaling, response to growth factors, inflammatory response or leucocyte migration (**Fig 5A and Suppl Table 4**). Additionally, considerable number of DEGs involved in cell differentiation and cell death were found to be upregulated in males upon tumor growth. Similar to males, sixteen processes were identified in females (**Fig 5B**), although all were classified under regulation of signaling pathways, immune and inflammatory processes as well as other processes such as cell death, transport regulation and positive regulation of angiogenesis. DEGs contributing to metabolic processes and cell proliferation and differentiation were not found in females, unlike in males (**Fig 5B**). Five specific processes were found to be common in both sexes. These included response to interferon-gamma, IL-1 signaling, leucocyte migration, inflammatory response and apoptosis (**Fig 5A and B**). However, the number and type of DEGs involved in each of these pathways were different between males and females **(Suppl Table 4 and 5)**. While no specific processes were identified for downregulated DEGs in males due to very few numbers of genes, downregulated DEGs in females were found to be associated with processes such as negative regulation of angiogenesis, locomotion, cell adhesion and cytoskeletal organization (**Fig 5C**). List of downregulated DEGs associated with each of the BPs in females in given in **Suppl. Table 6**. Additionally, we also identified specific transcription factors (TFs), ligands, peptides, growth factors, receptors, channels, enzymes, chemokines and cytokines that were differentially regulated in males and females upon tumor growth. Of all DEGs between both comparisons, we found 11 TFs, 19 ligands, peptides and growth factors, 24 channels and receptors, 65 enzymes and 14 chemokines/cytokines (**Fig 5D**). While the number of TFs were higher in male neurons upon tumor growth, female neurons had higher number of channels and receptors, enzymes and chemokines and cytokines, regulated post-tumor growth (**Fig 5E and F**). Taken together, these data indicate significant differences in regulation of genes, processes and pathways by tongue tumor, between males and females.

**Fig 5.**
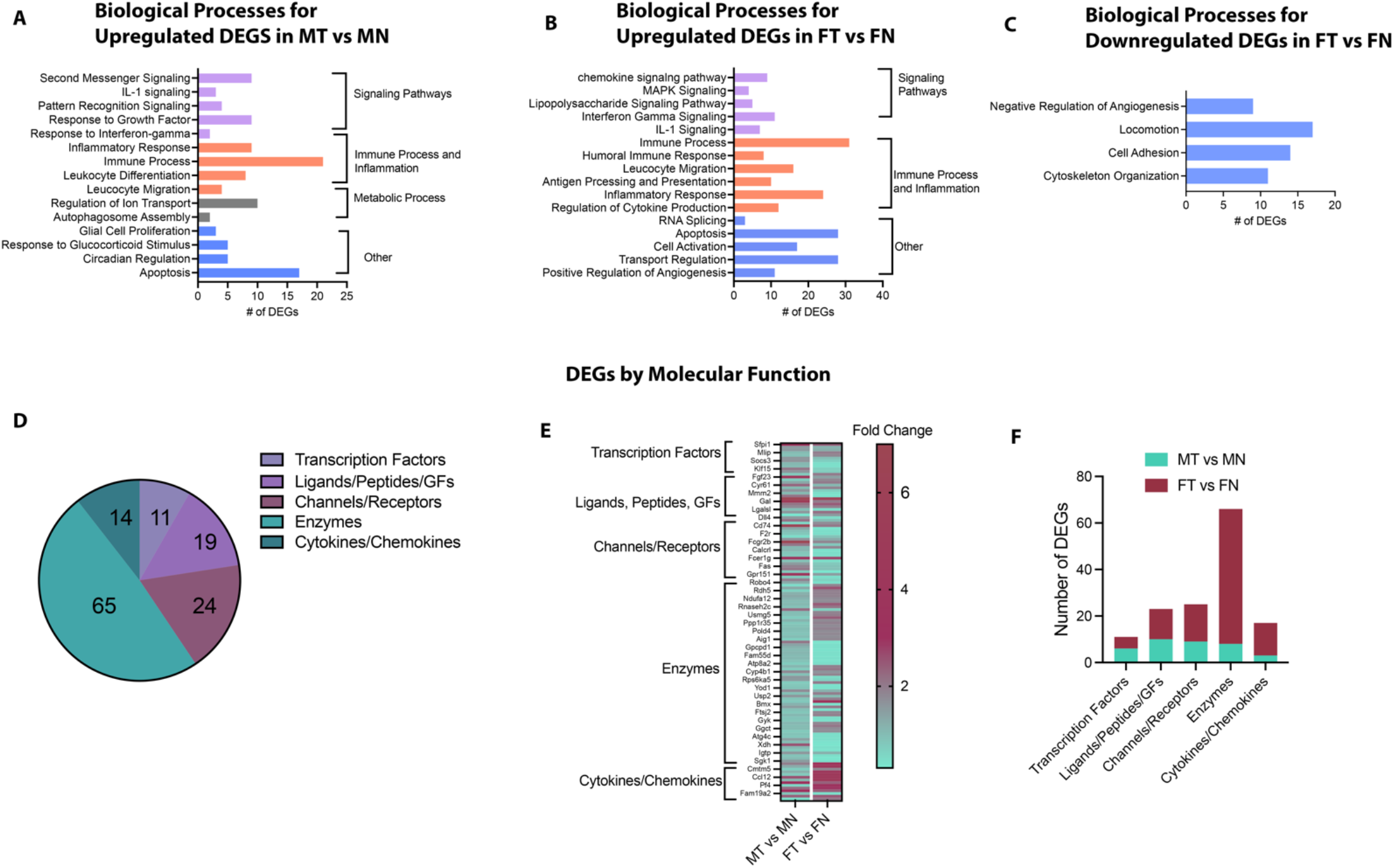
Biological Processes and Function of tongue-tumor controlled genes in males and females. Panther Pathway Analysis platform was used to elucidate biological processes from **A**. DEGs upregulated in MT vs MN, **B**. DEGs upregulated in FT vs FN and **C**. DEGs downregulated in FT vs FN. For each analysis, the number of DEGs associated with each biological process is plotted as bar graphs. **D**. Additional analyses was conducted to identify molecular function (MF) of all DEGs in males and females. Pie chart of the number of DEGs identified as transcription factors, ligands/receptors/growth factors (GFs), Channels/Receptors, Enzymes and Cytokines/Chemokines is shown. **E**. Heatmap of select DEGs for each of the molecular function indicate the differences in expression between MT vs MN and FT vs FN. Data plotted as fold change. **F**. Number of total DEGs obtained for each MF with MT vs MN and FT vs FN.

## Discussion

It is well established that chronic orofacial pain is sexually dimorphic with a higher prevalence in women than men for various pain conditions including temporomandibular joint disorders^50^, apical periodontitis^4^, oral mucositis^32,33,43^, burning mouth syndrome^31^ as well as oral cancer^42,46,47^. Yet, there is a large gap in knowledge about the mechanisms that lead to sex-dependent differences in orofacial pain. Moreover, sex-dependent global genomic changes in trigeminal neurons during disease is entirely unexplored. One prior study that reported changes in transcriptomic profiles of trigeminal ganglia in conditions of masseter muscle inflammation in rats, only included males ^51^. On the other hand, another study that investigated changes in gene expression of TG following neuropathic pain in males and females, did not evaluate the whole genome ^52^. Furthermore, both these studies employed whole ganglionic tissues that primarily represent non-neuronal population as recently confirmed by *Mecklenberg et al* ^7^. To address this drawback, a third study employed single-cell sequencing of human and mouse TG tissues to implicate genes and cell types in migraine. However, sequencing of TG tissue in this study only evaluated sex-differences under naïve conditions and genomic data obtained from mouse migraine models was not separated by sex^10^. Therefore, no study till date has investigated changes in transcriptomic profiles of the trigeminal sensory neurons during disease in both sexes. Furthermore, it is critical to study neurons-innervating specific tissues over all ganglionic neurons as it is indicated that type of sensory innervation is tissue-specific. For example, it has been reported that dental pulp innervating neuronal soma primarily are large diameter myelinated neurons unlike other tissues in the orofacial region such as the skin or mucosa^53,54^. Accordingly, percentage of TRPA1 expressing neurons are higher in the oral mucosal tissues than in dental pulp^54^. Similarly, we have shown that about 20% of tongue innervating sensory neurons are CGRP positive ^13^ whereas almost 50% of masseter muscle innervating neurons are CGRP positive^15^.

Therefore, in the current study, we explored changes in genomic profile of isolated sensory neurons innervating mouse tongue tissue and identified sex-dependent differences. We employed the approach of bulk-RNA sequencing of FACS sorted retro-labeled mouse tongue innervating TG neurons, to be able to concentrate neuronal population from whole ganglionic tissue as well as explore tissue-specific differences in neurons between sexes. Additionally, we utilized the preclinical mouse tongue cancer model as our disease model, as it has been shown that patients with tongue cancer are more commonly in pain than other oral cavity cancers and it is widely included in clinical and preclinical studies of oral cancer pain ^35-40^. However, till date, there is only a couple of studies investigating sexually dimorphic mechanism for tongue cancer pain ^42,46^. While these studies demonstrated the role of the immune response in sex-dependent differences in oral cancer pain, differential gene expression in sensory neurons upon tumor growth cannot be excluded, as we and others have reported that sensory neuronal activity is altered upon oral tumor growth^27,36,37 55-57^.

FACS sorting of lingual TG neurons was performed from normal and tumor-bearing animals for both sexes and our data indicated that female mice have higher number of tongue-innervating sensory neurons than males as observed by significant increased percentages of sorted cells from female samples than male samples. In further exploring this result, we found that female mice had higher percentage of TRPV1- and lower percentage of TRPV1+ neurons innervating the tongue than in male mice. However, recently *Scheff et al*, demonstrated no difference in capsaicin sensitivity in the tongue between males and females under naïve conditions ^56^, indicating that while the percentage of TRPV1+ neurons are different between the two sexes, the total number of TRPV1+ lingual neurons may not be different between males and females. Therefore, given that the total number of tongue-innervating neurons are higher in females, our data suggests that the total number of non-nociceptive sensory neurons in mouse tongue may be higher in females than in males. To our knowledge, this is the very first evidence of differences in number of neurons innervating orofacial tissues between sexes. Whether or not the increased innervation of non-nociceptive neurons contributes to sex-dependent pain observed in any of the chronic lingual diseased states is yet to be determined. Besides, whether this finding is specific to mice or exists in tongue tissues of higher order species such as human and non-human primates is unknown.

In analyzing RNA sequencing data between naïve male and female neurons, we found that several genes (i.e 30 genes) were exclusively expressed in females and not in males. This was in accordance to previous report showing increased DEGs in naïve female versus male neurons from whole mouse trigeminal ganglia^7^. Interestingly, many of these DEGs such as CSF1R (colony-stimulating factor receptor 1)^58,59^, C1qa (complement component 1 q) ^60-62^, Sh2b3 (lymphocyte adapter protein)^63^,Hhex (Hematopoetically expressed homeobox) ^64,65^ and Retnlg (Resistin-like gamma)^66^, are known to be primarily expressed in immune cells and have shown to play various roles in immune processes^58,62,67^, myeloid cell differentiation^64,65^, cytokine signaling^63^ and inflammation^66,68^. In fact, some of these genes such as Csf1R and C1qa are considered specific markers of mononuclear phagocytic system such as microglia, monocytes, macrophages and dendritic cells^58,59,61,62^. Csf1R in microglia and macrophages has been reported to be activated in neuropathic pain via its ligand Csf1 expressed in immune cells or in sensory neurons ^67^. Similarly, the contribution of microglia-expressed C1q proteins have also been reported in pain including orofacial pain^69-71^. Additionally, it has been shown that Csf1R exerts its action in a sex-specific manner by favoring a response in males compared to females^72^. However, for the first time, we report the expression of Csf1R in trigeminal sensory neurons, although its expression is specific to female neurons. Our immunohistochemical analyses of Csf1r and C1qa confirmed our RNA sequencing result and revealed that while majority of C1qa was expressed in non-TRPV1 expressing neurons, Csf1R was expressed in nociceptive and non-nociceptive neurons. Moreover, we also found that Csf1R signaling by its ligand Csf1 sensitizes TRPV1 responses in female sensory neurons but not in males indicating that at least some of the DEGs exclusively expressed in females may contribute to nociception. Interestingly, this effect was observed in WGA+ and WGA-cells suggesting the role of Csf1/Csf1R signaling in females may not be restricted to the tongue tissue. Collectively, these data indicate that perhaps female sensory neurons may be functionally different than male neurons innervating the mouse tongue.

Our analyses for gene expression changes post tongue-tumor growth revealed altered expression of several DEGs in both sexes. However, the number of DEGs in females was considerably higher than in males (83 DEGs in males vs 382 DEGs in females) suggesting that male neurons are more tightly regulated in spite of the tumor compared to female neurons. It would be interesting to investigate whether this result is specific to tongue tumors or even other lingual pathologies. Notably, while we observed similar pain behaviors (as measured by previously reported feeding behavior^27^) and tumor growth at the selected time point between both sexes (**Fig S1A and B**), there were only 18 genes that were commonly altered between males and females, indicating that the mechanisms of neuronal regulation in oral cancer may be considerably different between sexes, even if pain manifestation is similar. Clinically, association of gender differences in oral cancer pain has been varied. While, several studies report no association of gender and oral cancer pain^73-82^, one study reported that initial presence of pain may be more common in men^35^ and another study reported higher levels of function-related pain in men than women^47^. However, three other studies report worse pain scores for women than men.^83,84^. Pre-clinically, differences in pain behaviors appears to be varied as well and may depend on the tumor model used. For example, *Scheff et al* ^46^, showed that females had worse gnawing behavior in the 4NQO tongue cancer model compared to males whereas we observed no difference in feeding behavior in the current study with HSC3 tongue tumors (**Fig S1**). Nonetheless, these data indicate that it is crucial to understand the differences in mechanisms of oral cancer pain between males and females, to obtain useful insights about treatment options for both sexes.

Since the tumor developed in both sexes was from the same cell line (i.e HSC3), perhaps the differential regulation of neurons between males and females may be due to stark differences in the tumor microenvironment or alterations occurring at the ganglionic level upon tumor growth. Intraganglionic activation of macrophages ^85^ and satellite glial cells ^86,87^ has been reported in pain. While no studies have yet probed the contribution of these cells for oral cancer pain, a similar response within the TG can be expected that in turn would lead to changes in the neuronal soma. On the other hand, the peripheral tumor microenvironment consists of various cell types including keratinocytes, fibroblasts, endothelial cells, Schwann cells, and immune cells. Schwann cells ^88^ and immune cells^42,46^ have already been shown to contribute to tongue cancer pain and as mentioned above, immune cells even play a sex-dependent role in tongue cancer pain ^42,46^. Therefore, examining the role of these cell types in mediating global changes in the neurons of males and females would be crucial in better understanding of the impact of the tumor microenvironment in sexually dimorphic tongue cancer pain.

It is noteworthy though that despite only 18 DEGS common to both males and females, 5 biological processes (BPs) were found to be common between males and females post tumor growth. These included apoptosis, response to interferon gamma, inflammatory response, leucocyte migration and IL-1 signaling. Of the 18 common DEGs, four genes were associated with two of the common BPs; i.e inflammatory process and apoptosis. These four genes were Timp 1(tissue inhibitor of metallopeptidase 1) Gal (galanin), Chi3l3 (chitinase-like 3) and Cxcl10 (chemokine-ligand 10). Timp1 is a glycoprotein that is known to promote cell proliferation and anti-apoptosis; is implicated in cancer progression^89^; and negatively regulates matrix metalloproteinases and disintegrin-metalloproteinases (ADAMs)^90^. Interestingly, ADAM17 is implicated in oral cancer pain^91^. Accordingly, Timp1 has been shown to attenuate inflammatory pain in preclinical models^92^.

Galanin is a neuropeptide and is considered a potent modulator of inflammation by promoting cytokine production in immune cells^93,94^. It also has been shown to induce cell death in pheochromocytoma cells^95^. Importantly, the role of galanin in pain has been reported to have pro- and anti-nociceptive functions and whether or not galanin plays a role in peripheral nociceptive mechanisms is yet to be confirmed^96^. Interestingly, galanin release from sensory neurons have been shown to promote oral cancer progression^97^.

Chitinase-like proteins belong to the family of glycoside hydrolase and are involved in the regulation of the innate immune response^98^. Chi3l3 has been specifically reported to orchestrate recruitment of eosinophils in meningitis and autoimmune neuroinflammation^99,100^. However, its role in pain is not yet defined.

Cxcl10 is an important chemokine for inflammatory processes and its function in pain has been studied in several pain models including neuropathic pain and inflammatory pain^101-103^.

Aside from the above-mentioned four genes, many other genes were selectively altered in each sex, yet associated with the common BPs. For example, the cytokine, interleukin-1beta (IL1β) was specifically shown to be induced in female post tumor growth and was not expressed in males. The role of IL1β has been widely reported in pain, with the cytokine mostly produced by immune cells and other non-neuronal cells during injury^104,105^. However, because most of the studies have been conducted using male animals, neuronal IL1β has not been reported previously. To this end, a sex-specific role of IL1β in pain has not been studied till date. Interestingly, while induction of neuronal IL1β was only observed in females, downstream signaling of IL1 pathway was observed in both sexes post tumor growth indicating that perhaps IL1 β in males may be increased in the periphery or within the ganglia in non-neuronal cells as demonstrated in other injury models^104-107^.

Another example was expression of Hspa1a which encodes for heat-shock protein 70 (Hsp70). Our RNA sequencing data showed that while this gene was expressed in both sexes, it was specifically upregulated only in males post tumor growth. Interestingly, hsp70 has been demonstrated to have a protective role in pain during nerve damage^108,109^, migraine ^110^ and opioid-induced hyperalgesia^111^.Therefore, upregulation of Hsp70 in males post-tumor growth might indicate that male neurons might express an endogenous feedback mechanism to suppress pain that may be lacking in females.

In addition to the common BPs, sex-selective BPs were identified post-tumor growth. Noteworthy female-selective BPs including cytokine/chemokine production and signaling as well as angiogenesis. The contribution of various cytokine/chemokines in nociception and inflammation^112-114^ is widely established and our data showed that the number of cytokine/chemokines altered in female neurons was higher than in males (i.e. 14 in females versus 3 in males). This is an intriguing yet not a surprising observation as it aligns with the finding that female neurons express immune cell markers as described above.

Female DEGs were also associated with angiogenesis. Interestingly, genes associated with positive regulation of angiogenesis (e.g *Gadd45a, Dcn, Ngp, cxcl10, Hmox1,ccl11,Pgf etc)* were upregulated upon tumor growth and those associated with negative regulation of angiogenesis (e.g. *Cav2, Ptprm, Mmrn2, Xdh*, etc) were downregulated, indicating that sensory neuro-vascular interaction may be sex-specific.

In contrast, male-specific BPs included growth factor signaling and ion transport. One of the top genes upregulated in males upon tumor growth; fibroblast growth factor 23 (Fgf23) is reported to be associated to both of these processes. It is not only a growth factor that induces downstream signaling via its receptors ^115^ but also controls phosphate homeostasis ^116^. Furthermore, it is also considered a bone-derived hormone that is regulated by inflammation and associated with bone pain ^117,118^.

Taken together, our data points to significant differences in the regulation of lingual sensory neurons in males and females upon tongue tumor growth. The current study is significant as it is the first to comprehensively characterize the genomic profile of tongue-innervating neurons under naïve and tumor-bearing conditions. Our data lays the foundation for future investigations to identify potential sex-specific targets in sensory neurons and study their functional and mechanistic contribution in tumor-induced pain. Of interest are the DEGs belonging to the class of transcription factors in males and enzymes and cytokine/chemokines in females as the total number of DEGs in these classes were higher in the respective sexes. One limitation of the study include use of one cell-line to induce the tongue tumor and at this point, the study cannot confirm which DEGs and processes are common across tumors induced by different OSCC cell lines. We also note that the study was conducted using the xenograft model with athymic mice that lack the response of T lymphocytes. While this model allows to explore the regulation of sensory neurons by human tumors, it excludes the involvement of T cells in tumor-induced pain. Therefore, identifying DEGs in both sexes that are common to the xenograft as well as the syngeneic model will be important in selecting relevant targets for functional studies. Besides, it would be useful to delineate the temporal effect of the tumor on neuronal genes and processes to further gain insight into sex-specific regulation of sensory neurons in oral cancer.

## Materials and Methods

### Animals

Six-to eight-week-old adult Balb/c male and female athymic nude mice (Jackson Labs, Bar Harbor, ME, USA) were used for all experiments. All animal handling and procedures were performed according to approved UTHSCSA IACUC protocols and conformed to the guidelines of International Association for the Study of Pain (IASP). Animals were housed in the UTHSCSA laboratory of Animal Resources (LAR) for at least 4 days prior to start of experiments.

### In Vivo Orthotopic Xenograft Tongue Tumor Model

#### Using human oral squamous cell carcinoma cell lines

HSC3, tongue tumors were induced in mice as described by us previously^27,36,37^. Animals were anesthetized with isoflurane inhalation and 50ul of 3.5 x 10^5 HSC3 cells were injected unilaterally in the ventral side of the tongues using insulin syringes. Animals were then allowed to recover in their cages. Animals were used for experiments at day 15 post-cell inoculation.

### Retro-labeling of Tongue-Innervating TG neurons

Tongue-innervated sensory neurons were labeled as described by us previously^13^. Briefly, animals were anesthetized with isoflurane inhalation and 10uls of 1% wheat germ agglutinin (WGA)-AF488 (Promega), diluted in 1%DMSO, was injected bilaterally twice in tongue of each animal. The first injection was given in the superficial epithelial layer and 4 hours later, a second injection was given in the deeper muscular layers of the tongue. Animals were allowed to recover in their cages for 2 days before harvesting trigeminal ganglia (TG) tissues.

### Preparation of single-cell suspension and flow sorting

#### Tongue-innervating sensory neurons were isolated from 4 groups

Male normal (MN), Female normal (FN), Male tumor (MT) and Female tumor (FT). Neurons were isolated by preparation of single-cell suspensions of TG tissues followed by flow sorting of WGA+ cells. Each sample was prepared by pooling 4 TGs from 2 animals for normal groups and 4 animals for tumor-bearing groups as only ipsilateral TGs were collected from this group. A total of 3 samples per group was prepared. TG tissues were dissected and collected in cold 1X HBSS buffer, washed three times with 1X HBSS and incubated with 5ul of 50ng/ml dispase (type 2, Sigma) and 75ul of 2.5mg/ml liberase (Roche) for 60 mins at 37°C for enzymatic digestion. Following this, tissues were centrifuged at 2min at 1000rpm and washed with 5mls of DMEM containing 5% FBS and resuspended in 1.5ml DMEM with 5% FBS and triturated with a Pasteur pipette to breakdown the tissues and prepare a homogenous solution. The solution was then strained with a 100um strainer to remove all debris, supernatant collected in an eppendorf tube and subjected to flow sorting. Flow sorting was performed using FACSAria III (BD Biosciences; San Jose, CA) using 130 μm nozzle. Consecutive gates were used to isolate WGA-AF488 labeled TG neurons. First, debris was excluded by forward scatter area (FSC-A) and side scatter area (SSC-A) gating. Second, duplets and clumps were excluded by side scatter width (SSC-W) and side scatter area gate (SSC-A) gate. Third, WGA-AF488+ bright cells were gated compared to unstained TG control and sorted directly to RLT buffer (Qiagen) containing 1% 2-mercaptoethanol (Sigma) to be able to use for RNA extraction.

### RNA isolation

RNA was extracted using Qiagen RNeasy micro kit (Qiagen) with on column DNase I digestion according to manufacturer’s instructions. RNA quantity was evaluated using Agilent 2100 Bioanalyzer RNA 6000 Nano chip (Agilent Technologies, Santa Clara, CA).

### Bulk RNA sequencing

Bulk-RNA sequencing of samples was conducted at the UTHSCSA Genome Sequencing Facility. RNA integrity was determined using Fragment Analyzer (Agilent, Santa Clara, CA) prior to library preparation. All samples were ensured to have RIN values >6.5 to proceed with library preparations. RIN values for each sample is listed in **Supplementary Table 1**. RNA-seq libraries were prepared according to SMART-seq2 protocol ^119^, with the following modifications: PCR preamplification to 10-12 cycles, two rounds beads cleanup with 1:1 ratio after cDNA synthesis, and 0.6-0.8 dual beads cleanup for Nextera XT DNA-seq library purification. RNA-seq libraries were sequenced using Illumina HiSeq 3000 system (Illumina, San Diego, CA) with 50bp single-read sequencing module. Upon sequencing completion, short read sequences from RNAseq were first aligned to UCSC mm9 genome build using TopHat2 aligner and then quantified for gene expression by HTSeq to obtain raw read counts per gene, and then converted to RPKM (Read Per Kilobase of gene length per Million reads of the library).

### Analyses of RNA sequencing Data

Sequencing data was analyzed as previously described^120^. Briefly, differential expression analysis was performed using DESeq ^121^ algorithm (R/Bioconductor) to estimate the differential expression in read counts and their statistical significance. Significantly differentially expressed genes were selected based on following criterion: 1) RPKM > 5 in one of the comparison group, 2) fold-change > 1.5, and 3) differential expression p-value < 0.05 along with False-discovery rate (FDR) for each gene. When comparing female tumor samples with male tumor samples, all genes coded in sex-chromosomes (chrX and chrY) were excluded. Functional assessment of DEGs were performed by using over-representation statistic (Fisher’s Exact test) for Gene Ontology using Panther platform.

### Immunohistochemistry

Protocol for immunostaining is described previously by us^13,36^. Mice were anesthetized with ketamine (75mg/kg)/ dexmedotomidine (1mg/kg) solution and perfused with 4% paraformaldehyde (PFA) and trigeminal ganglia was dissected bilaterally from each mouse. Tissues were post-fixed in PFA, washed in 0.1M phosphate buffer and cryosectioned in Neg-50 (Richard Allan, Kalamazoo, MI, USA) at 20uM thickness. Sections were then subjected to immunostaining with specific primary antibodies as described in **Table 1**. Donkey anti-secondary antibodies were purchased from Molecular Probes, Eugene, OR, USA and were used at a dilution of 1:200 for all experiments. Following staining, tissue sections were mounted in Vectashield and imaged using Nikon Eclipse 90i microscope equipped with a C1si laser scanning confocal imaging system. Z-stack images were acquired of the V3 region of TGs from at least 2 animals per group and a total of at least 6-10 images were taken for each antibody combination per group. All images were obtained with a 20x objective at fixed acquisition parameters across all groups and were unaltered from that initially taken. Laser gain settings were determined such that no-primary control did not show any positive staining. Quantitation was achieved using Adobe Photoshop 2023 for number of neurons above threshold, in the V3 region of the TG tissue in each image. A total of 200-500 neurons per analyses were counted.

**Table 1.** List of Primary Antibodies used for Immunohistochemistry

| Marker | Company | Catalog Number | Dilution |
| --- | --- | --- | --- |
| TRPV1 | Neuromics, Edina, MN | GP14100 | 1:200 |
| CSF1R | ThermoFisher Scientific, Waltham, MA, USA | 14-1152-85 | 1:50 |
| C1qa | Novus Biologicals, Centennial, CO, USA | NB200-539 | 1:50 |
| IL1-Beta | Novus Biologicals | NB600-633 | 1:50 |
| HSP70 | Novus Biologicals | NBP1-77456SS | 1:50 |
| **Table 1. List of Primary Antibodies used for Immunohistochemistry** |  |  |  |

### Primary Culture of TG neurons

Mouse TG neurons were cultured for calcium imaging experiments. Naïve male and female mice were injected with WGA as described above, bilaterally in the tongue and 2 days later, TGs were dissected for neuronal cultures as described previously^122^. Tissues were washed in HBSS and then dissociated with 1mg/ml of collagenase and dispase (Roche, Indianapolis, IN, USA) for 45 mins at 37 degrees. Following washing out the enzymes, cultures were plated on poly-D-lysine/laminin coated glass coverslips (BD Biosciences, San Jose, CA, USA) and grown overnight in either DMEM media supplemented with glutamine, penicillin/streptomycin and 2% Fetal bovine serum (for vehicle group) or the same media with 100ng/ml recombinant colony-stimulating factor 1 (Csf1, Cell Signaling, Boston, MA, USA). Csf1 stock was diluted in saline.

### Calcium Imaging

Calcium accumulations of cultured sensory neurons was determined by fluorescence imaging as described previously 122. TG cultured were incubated with calcium-sensitive dye, Fura-2 AM (2μm; Molecular Probes, Carlsbad, CA, USA) in Hanks modified buffer. Imaging was performed with a Nikon TE 2000U microscope fitted with a ×40/1.35 NA Fluor objective. Following selection of small-to-medium neurons and baseline measurements, cells were applied with 15nM capsaicin (CAP) to record TRPV1 responses. The net changes in calcium influx were calculated by subtracting the average baseline intracellular calcium [Ca2+]i level from the peak [Ca2+]i value achieved after exposure to CAP.

### Feeding Behavior

Feeding behavior was performed as previously described by us^27,36^. Briefly, animals were food deprived for 18 hours in their cages with bedding, following which, animals were placed in individual cages without bedding to acclimate for 30min to an hour. Premeasured standard chow was then provided to the animals to feed freely for 1hr. At the end of the hour, food intake was measured by calculating the difference (in g) from the initial amount and the remaining amount of food. Baseline measurements and food intake at day 15 post HSC3 cell inoculation was determined for male and female mice.

### Tumor Volume Measurement

Tumor volumes were measured at day 15 post HSC3 cell inoculation in male and female mice as described previously^36^. Animals were anesthetized with 2% isoflurane inhalation and tongue tumors were measured by determining length, width and height of the tumors using calipers. Tumor volumes were calculated by the formula π/6 (length x width x height) and presented as mm3.

### Statistical Analyses

Sample sizes were calculated using G-power application to obtain 80% power at a two-sided tail with α error probability of 0.05. All animals were allocated to groups using simple randomization. All statistical analyses and graphical representations were performed in GraphPad Prism 9.0. Data are presented as mean ± standard error of the mean (SEM) or as heatmaps and volcano plots for RNA sequencing data. Statistical significance was determined using either Paired Student T-test, Unpaired Student’s T -test, One-way ANOVA or Two-way ANOVA with Sidak’s post-hoc test and p<0.05.

## Acknowledgements

Study was supported by funds provided by NIH R01DE027223 (SR) and R01DE029187 (SR) and by the National Institute of Dental and Craniofacial Research. RNA sequencing data was generated in the Genome Sequencing Facility, which is supported by UT Health San Antonio, NIH-NCI P30 CA054174 (Cancer Center at UT Health San Antonio) and NIH Shared Instrument grant S10OD030311, and CPRIT Core Facility Award (RP220662).

## Author Contributions

T.I. performed animal injections for tumor growth, WGA injections and conducted all immunohistochemical experiments, P.W. performed TG dissections and single cell suspensions for Flow sorting. A.T. performed flow sorting and RNA extractions, L.Z and K.W prepared cDNA libraries and performed RNA sequencing. Y.C and W.I analyzed RNA sequencing data along with S.R. J.M. (Murillo) performed behavior assays. J.M. (Merlo) and C.F.M. performed calcium imaging experiments. S.R. conceptualized, designed and analyzed experiments, as well as wrote and edited the manuscript. All authors edited the manuscript.

## Data Availability Statement

Data that support presented findings are available as figures, materials and methods as well as supplementary tables. Additionally, all RNA-seq data is uploaded to NCBI (Accession Number GSE220448) and released as soon as manuscript is accepted for publication.

