## supplemental information for "Sex-dependent Differences in the Genomic Profile of Lingual Sensory Neurons in Naïve and Tongue-Tumor Bearing Mice"

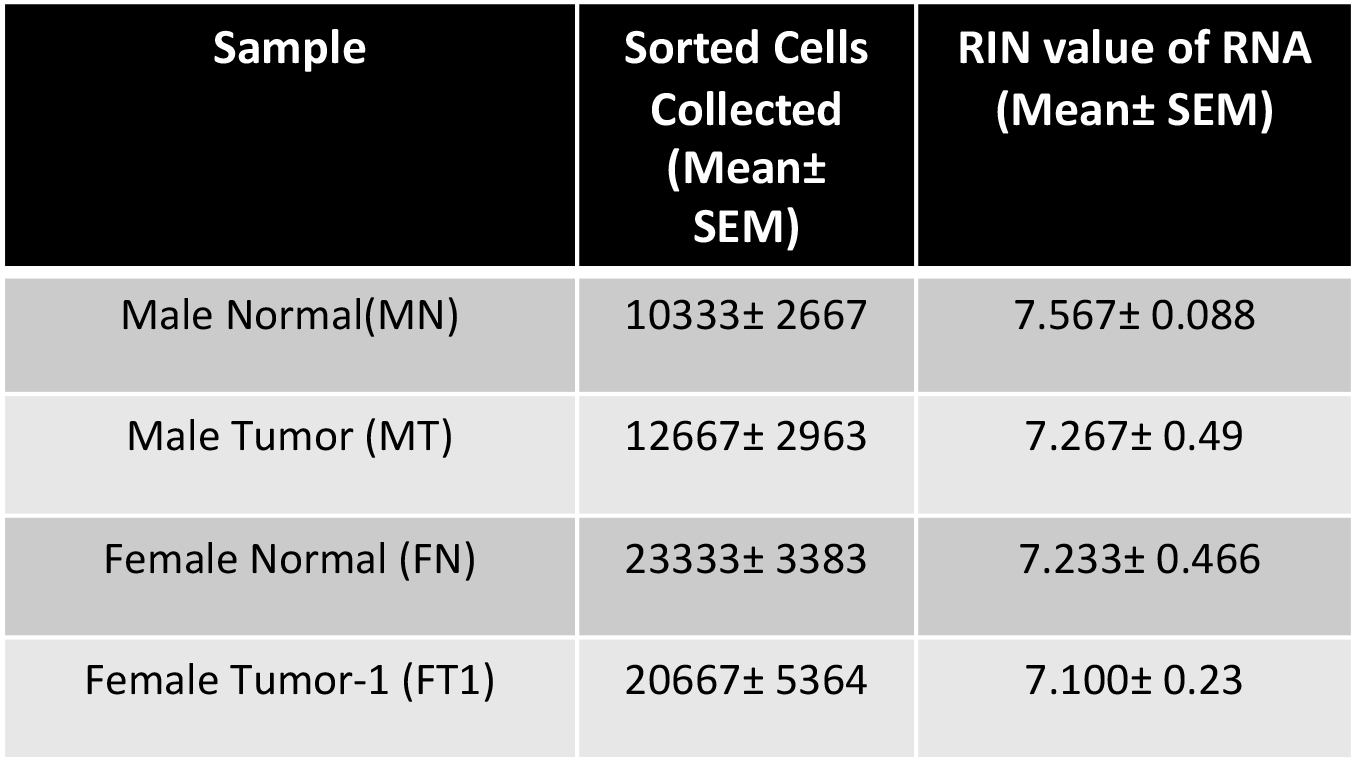

**Supplementary Table 1.** Number of sorted cells and RIN values of RNA of all samples within four groups. Data represented as mean ± SEM.MN: Male Normal. MT: Male Tumor. FN: Female Normal, FT: Female Tumor. N=3 per group.

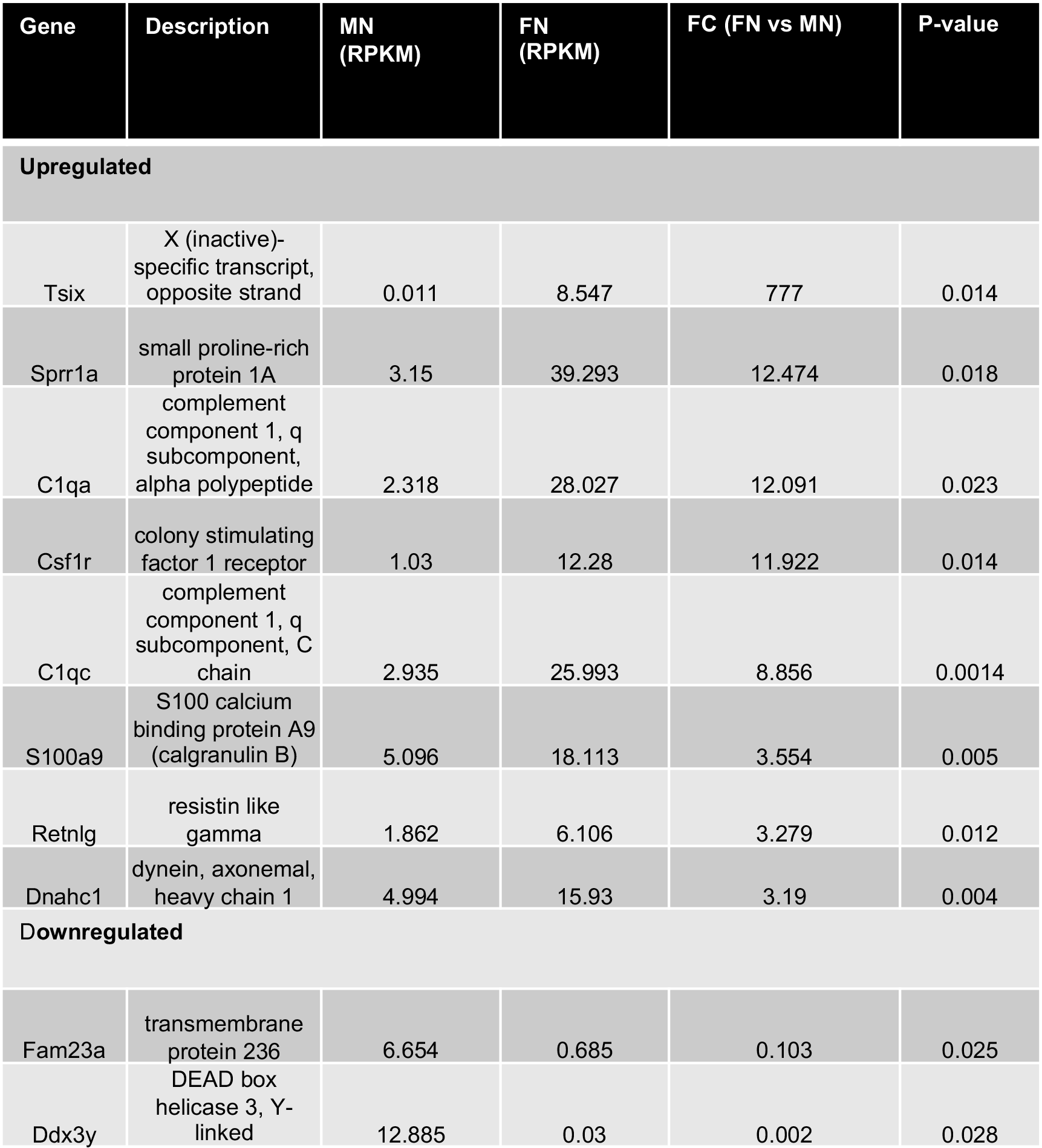

**Supplementary Table 2.** Top 10 DEGs of FN vs MN.

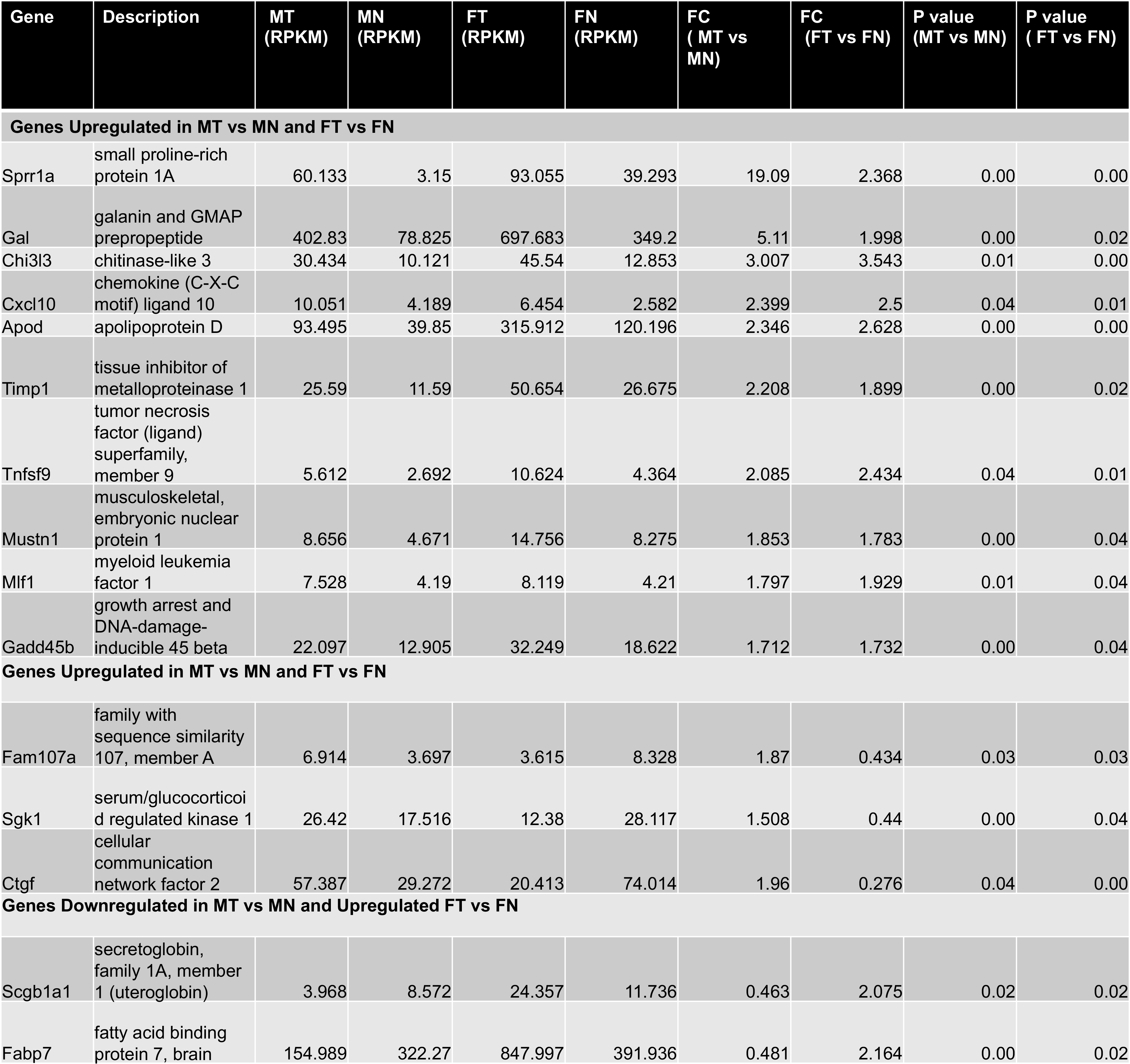

**Supplementary Table 3.** DEGs common to MT vs MN and FT vs FN

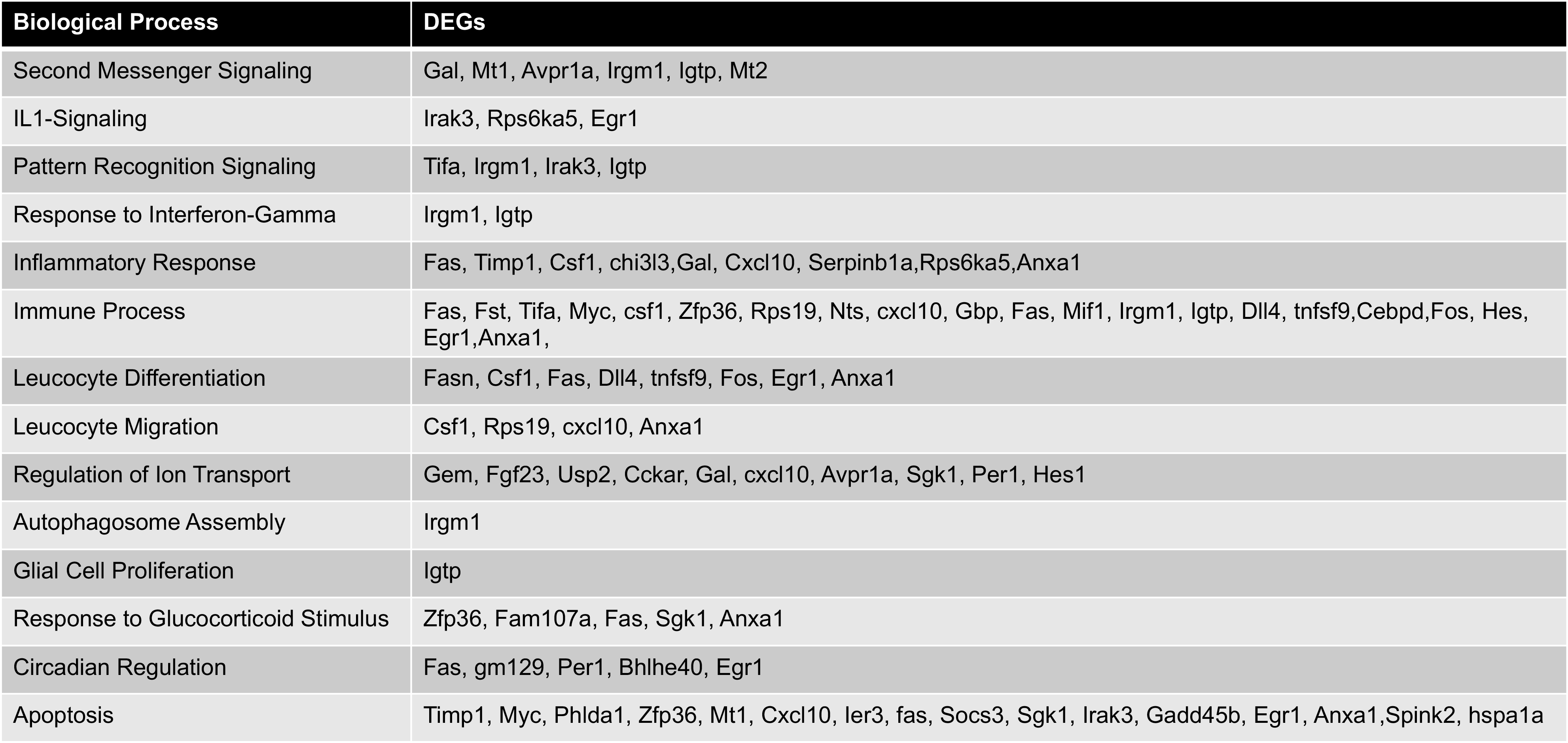

**Supplementary Table 4.** Biological Processes associated with DEGs upregulated in MT vs MN

**Supplementary Table 5.** Biological Processes associated with DEGs upregulated in FT vs FN

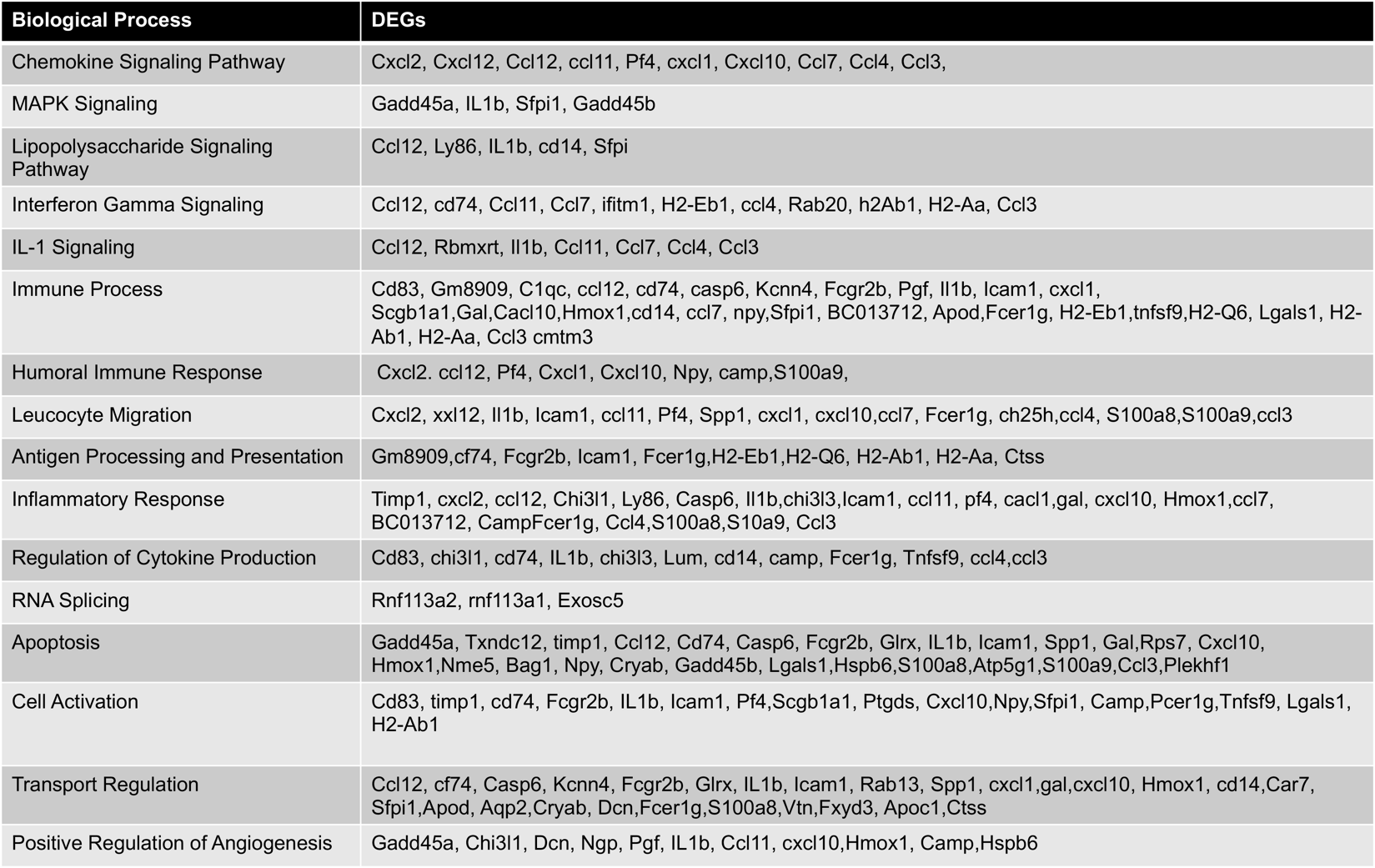

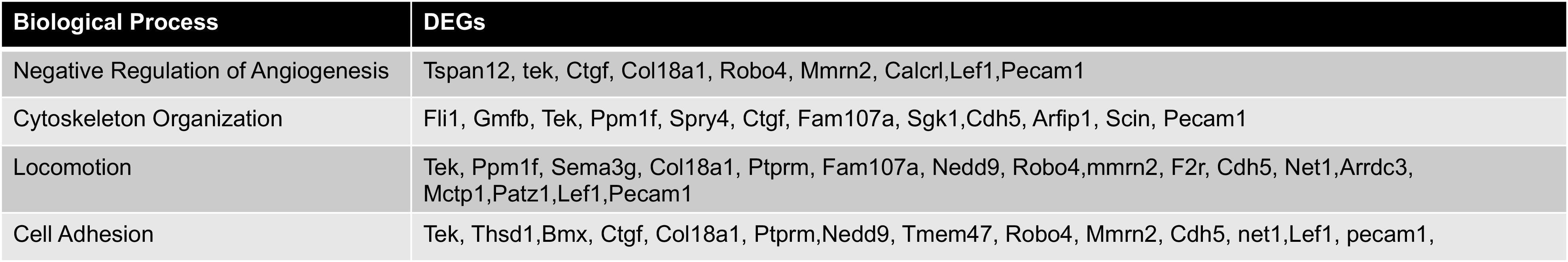

**Supplementary Table 6.** Biological Processes associated with DEGs downregulated in FT vs FN

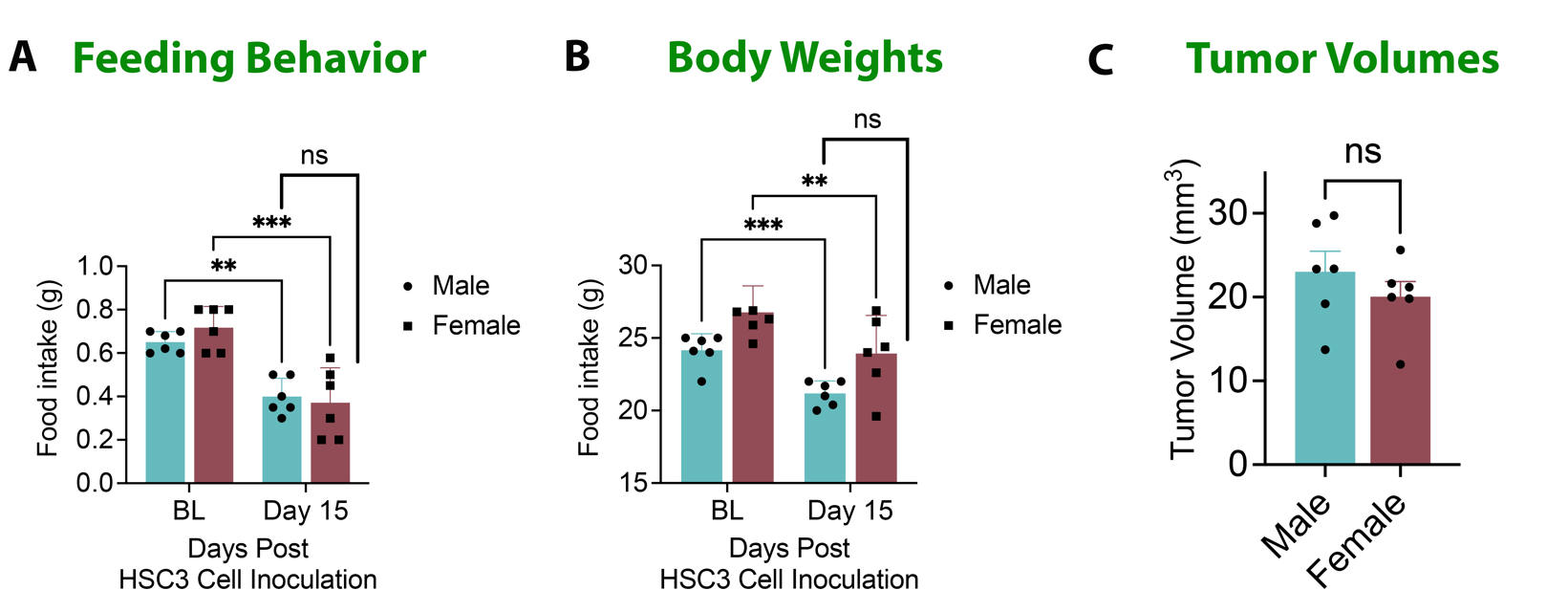

**Supplementary Figure 1. Pain Behavior post-HSC3 Tongue Tumor Growth in Males and Females.** Male and Female mice were injected with 3.5x10^5 HSC3 cells in the tongue. **A**. Feeding Behavior was determined at baseline (BL) and at day 15 post cell-inoculation. N=6. Data are presented as mean ± SEM and analyzed by one-way ANOVA with Sidak’s post-hoc test p<0.05. **B.** Body weights were determined at baseline (BL) and at day 15 post cell-inoculation. N=6. Data are presented as mean ± SEM and analyzed by one-way ANOVA with Sidak’s post-hoc test p<0.05. **C**. Tumor Volumes were measured at day 15 post cell-inoculation. N=6. Data are presented as mean ± SEM and analyzed by Unpaired Student’s T-test at p<0.05.
